# A novel regulatory element upstream of *TNFAIP3* promoter may be influenced by SLE risk variants

**DOI:** 10.1101/2020.09.30.320119

**Authors:** Ajay Nair, Archana S Rao

## Abstract

Almost all of Genome Wide Association Studies (GWAS) hits come from non-coding DNA elements. Data from chromatin interaction analyses suggest a long-range interaction with a putative enhancer upstream of *TNFAIP3*. Disrupting the enhancer may impair *TNFAIP3* expression and enhance SLE risk. Two variants, rs10499197 and rs58905141 carried on the SLE risk haplotype are situated near this enhancer and could affect its function. Cloning the regulatory region surrounding rs10499197 into an expression plasmid containing a CRISPR-Cas9 backbone, and then performing a genome editing assay, we found that the variant is located near an enhancer. And any changes to the SNP region might impair enhancer and its ability to regulate *TNFAIP3* expression.

## Introduction

Completion of the human genome sequence has enabled unparalleled exploration of the human genome. Availability of dense SNP panels with high-throughput, cost-effective genotyping and sequencing platforms, aid in performing unbiased high-resolution genome-wide surveys needed to identify DNA segments that harbor complex disease causal variants.

### Understanding SLE, Why it matters

Systemic Lupus Erythematosus (SLE) is an autoimmune disease characterized by dysregulated interferon response and loss of self-tolerance to cellular antigens. Autoantibody production leads to immune complex formation resulting in wide scale inflammation and multiple organ failure. SLE afflicts women at a rate nine times that of men (Chen et al., 2017; Deng and Tsao, 2017). The precise etiology of SLE is complex and poorly defined, requiring the interplay between inherited common risk alleles, genetic predisposition and environmental triggers.

### TNFAIP3, a key SLE-risk locus

Currently, there are more than 90 robust SLE susceptibility loci (Mohan and Putterman, 2015;Chen et al., 2017;Deng and Tsao, 2017). Testing association of 311,238 genotyped SNPs in 431 unrelated SLE cases and 2155 controls, three regions met a specified threshold for genome-wide significance: the previously defined *HLA, IRF5* regions and a novel SLE locus at chromosome 6q23 within the *TNFAIP3* gene (Dieude P. et al, 2010). *Tumor necrosis factor alpha-inducible protein 3* (*TNFAIP3*) is a prominent risk locus identified in many autoimmune diseases including SLE. *TNFAIP3* encodes the ubiquitin-editing enzyme, A20 that negatively regulates Nuclear Factor kappa-B (NF-κB) signaling to attenuate pro-inflammatory immune responses and protect against the damaging effects of chronic inflammation (Han et al., 2009; Cai et al., 2010).

Human GWAS of autoimmune diseases have identified single nucleotide polymorphisms (SNPs) on the *TNFAIP3* locus that encodes A20. However, due to strong linkage disequilibrium, GWAS cannot distinguish causal from benign SNPs. Our group previously identified several potential causal SNPs within the *TNFAIP3* gene locus associated with SLE. In this study, we aim to briefly evaluate the function of a novel regulatory element located upstream of the *TNFAIP3* promoter.

## Materials and Methods

### Antibodies and cell lines

Jurkat, Human embryonic kidney cells 293 (HEK293T) and THP-1 were purchased from ATCC. Epstein Barr Virus (EBV)-transformed B cell lines were obtained from the Lupus Family Registry and Repository (OMRF) with IRB approval. EBV cell lines were selected using genotype data corresponding to the rs10499197 variant. Genotypes were verified by Sanger sequencing. Cell lines were maintained in RPMI 1640 medium supplemented with 10% FBS, 1X penicillin-streptomycin antibiotic mixture (Atlanta Biologicals Inc., GA), 2mM L-glutamine (Lonza™ BioWhittaker™, Walkersville, MD), and 55μM ß-mercaptoethanol. The following antibodies were used in the study: anti-GAPDH (Cell Signaling Inc., Danvers, MA); anti-A20 (GeneTex Inc., Atlanta, GA); anti-Histone H3 (acetyl K27) (Abcam, Cambridge, MA); normal Rabbit IgG (EMD Millipore, Burlington, MA). All stock laboratory chemicals were from Sigma Aldrich(St.Louis,MO).

### Western Blotting

Two million EBV-transformed B-cells were pelleted and washed in cold PBS and lysed with RIPA lysis buffer (25 mM Tris-HCl, 150 mM NaCl, 5 mM EDTA, pH 8, 1% Triton X-100, 0.1% SDS, 0.5% sodium deoxycholate) containing proteases (EMD Millipore, Burlington, MA) and Halt phosphatase inhibitors (Pierce, Rockford, IL). Cells were lysed for 15 minutes on ice followed by syringe lysis with a 27g needle then cleared by centrifugation at max speed for 20 minutes. Protein concentrations were determined by Qubit assay (Life Technologies, Carlsbad, CA). Proteins were denatured in 2X SDS loading buffer by heating to 100 °C for 5-10 min. Proteins were eluted by SDS-PAGE and analyzed by immunoblotting as indicated.

### HiChIP Assay

A modified version of a previously published HiChIP method was used (Mumbach et al., 2016). Briefly, EBV-transformed B cells were crosslinked with 1% formaldehyde for 10 min at room temperature, then intact nuclei were isolated and DNA digested using MboI restriction enzyme (New England BioLabs, Inc., Ipswich, MA) for 4 hours at 37 °C. The ends of the DNA fragments were filled and labeled with dCTP, dGTP, dTTP, and biotin-dATP. Proximity ligation was performed at room temperature overnight, and then the ligated chromatin was fragmented using a Covaris S2 sonicator (Woburn, MA) and previously described parameters (Mumbach et al., 2016). The fragmented crosslinked protein/DNA complex was immunoprecipitated using an anti-H3K27acetylated antibody, and then purified by immunomagnetic beads (Magna ChIP™ Protein A+G Magnetic Beads, EMD Millipore Corp, USA). The number of cells in each sample was used to determine the amount of MboI enzyme, immunoprecipitating antibody, and immunomagnetic beads used (Mumbach et al. Nat Methods 2016). DNA was eluted from the beads by incubating at 65 °C for 4 hours with 5% Proteinase K and purified using spin column (DNA Clean & Concentrator™, Zymo Research, Irvine, CA). Purified DNA concentration was quantified by Qubit High Sensitivity Assay Kit (Thermo Fisher Scientific). The yields were used to determine the tagmentation enzyme amount and PCR cycle number. Library construction required a minimum of 2 ng DNA. Biotin-labeled DNA fragments (Thermo Fisher Scientific) were further immunoprecipitated by Streptavidin M-280 Dynabeads (Thermo Fisher Scientific). Finally, the HiChIP library was generated on the streptavidin beads using Nextera DNA Library Prep Kit (Illumina Inc., USA). The final PCR product was purified by two-sided size selection using Ampure XP beads (Beckman Coulter, Brea, CA) to capture DNA fragments between 300 bp and 700 bp. *Library quality assessment and sequencing:* Several quality controls were performed on the HiChIP library. First, the Qubit High Sensitivity Assay Kit was used to quantify DNA concentration. Secondly, the integrity and quality of the library was assessed by Agilent TapeStation 2200 bioanalyzer (Santa Clara, CA). A library was considered good if it had a clear DNA peak at 300-700 bp. Several HiChIP libraries were pooled together and quantified using the KAPA Biosystem library quantification qPCR kit (KAPA Biosystems, Wilmington, MA). The qPCR was performed on a Roche LightCycler 480 system using primers specific to the HiSeq flowcell. Using the template size determined from the Qubit, the pools were diluted to 2 nM and re-quantified using qPCR to confirm the starting concentration. The pooled samples were denatured for 5 min with 0.1N NaOH, and further diluted to a loading concentration of 1.8 pM. Libraries were sequenced on the Illumina NextSeq 500 sequencer on a 150-cycle paired-end Mid flowcell. *Analysis pipeline:* HiChIP raw reads (fastq files) were aligned to the hg19 human reference genome using HiC-Pro [Servant, et al., 2015]. Aligned data were processed and analyzed through hichipper pipeline [hichipper MS]. We used MAC2 for anchor calling in the pipeline based on ChIP-enriched regions. Loops were derived from the linked paired-end reads that overlap with anchors. A good quality sample has a minimum of 15% intrachromosomal interactions.

### CRISPR-Cas9 Genome Editing

A CRISPR-Cas9 system from reagent design, construction, validation and cell line expansion was developed using the established protocol (Ran F.A. et al., 2013). Custom guide RNA (gRNA) pairs for a 500bp sequence around the rs10499197 (T/G) SNP region, as well as genotyping primers, were designed *in silico* via the CRISPR Design Tool (http://tools.genome-engineering.org). gRNA guide sequences were cloned into pSpCas9(BB)-2A-GFP (PX458) expression plasmid bearing both gRNA scaffold backbone (BB) and Cas9, pSpCas9(BB). Completed and sequence-verified pSpCas9 (gRNA) plasmids and optional repair templates for facilitating HDR were then transfected into HEK293T cells and assayed for their ability to mediate targeted cleavage. Finally, transfected cells were clonally expanded to derive isogenic cell lines with defined mutations. Chimeric DNA generated was extracted and cleaned using a QIAGEN DNA purification kit (QIAGEN Inc., Valencia, CA). DNA was then amplified by PCR and subjected to Sanger sequencing to decode the clones.

## Results

### Epigenomic analyses revealed a putative enhancer upstream of the *TNFAIP3* promoter

Dynamic organization of chromatin in three-dimensional space, specifically enhancer-gene promotor looping, plays a crucial role in regulating gene transcription. To investigate the threedimensional organization of the TNFAIP3 locus, we performed highly integrative chromatin immunoprecipitation (HiChIP) analysis on chromatin obtained from Epstein Barr virus-transformed B cells (EBV) using antibodies against the epigenetic mark of an active enhancer, acetylation of Histone 3 lysine 27 (H3K27ac). We used hichipper and DNAlandscaper (Lareau and Aryee, 2018) to visualize, in two dimensions, several long-range enhancer-promotor interactions in and around the *TNFAIP3* locus (**Figure 1**). Notably, our H3K27ac HiChIP data revealed a prominent long-range looping event between the *TNFAIP3* promoter and a region ~55 kb upstream that is enriched with epigenetic marks of an enhancer (H3K4me1, H3K27ac) and clusters of transcription factor binding sites defined by ENCODE ChIP-seq data. The putative enhancer also overlapped SLE risk variants rs10499197 and rs58905141 located on the most 5’ end of the SLE-associated ~109 kb risk haplotype (**Figure 1**). Collectively these results suggest that the putative enhancer located ~55 kb upstream of *TNFAIP3* may play an important role in modulating *TNFAIP3* expression during SLE pathogenesis.

**Figure1.**
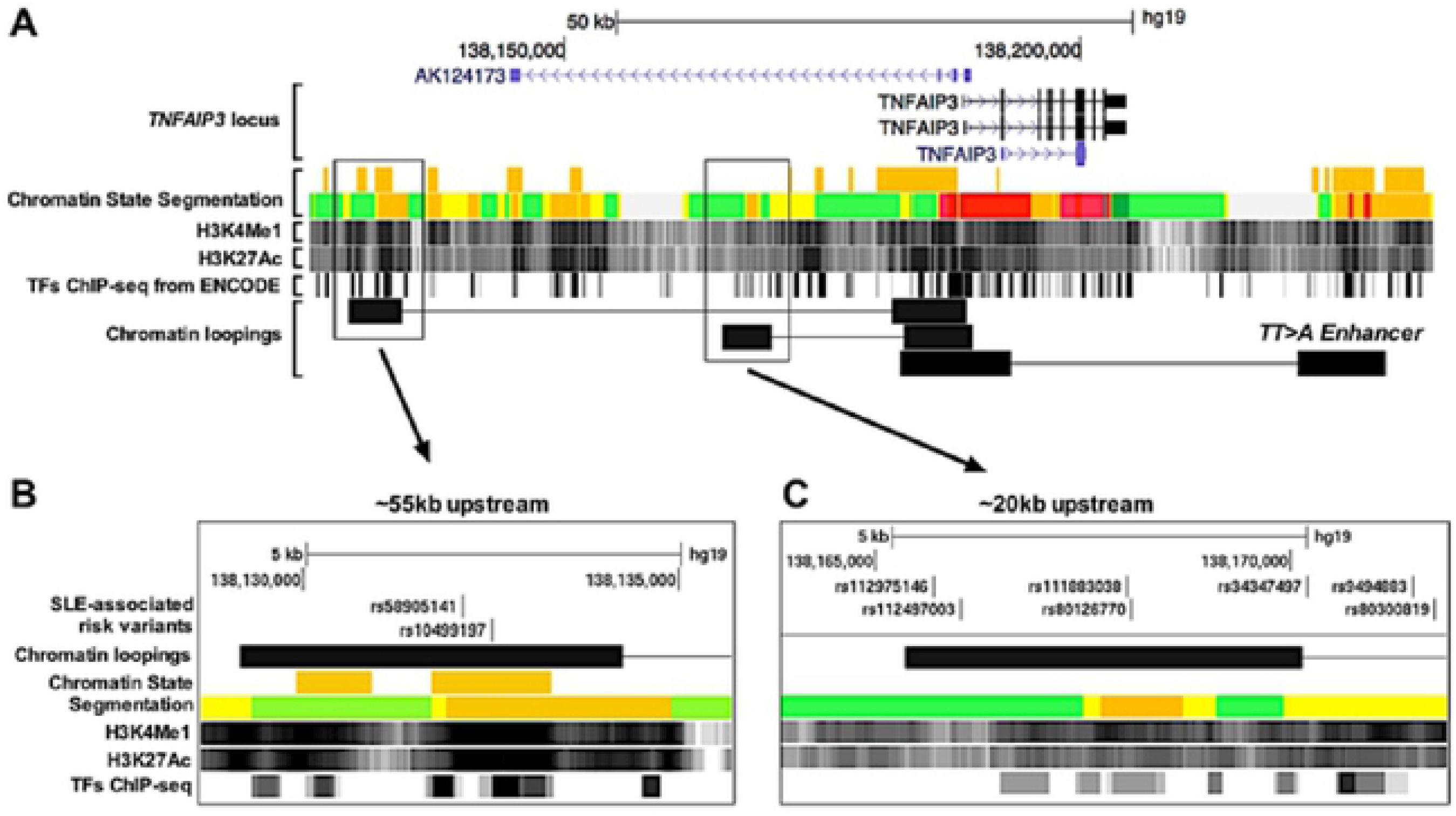
Location of SNP in putative enhancer region. Two risk variants locate in putative enhancer region. A. The relative locations of three most strong DNA loopings mediated by Pol2 in EBV cell lines. B, C. A zoom in view of two upstream interaction regions. The ENCODE Integrated Regulation super-tracks contain: Chromatin State Segmentation, H3K4Me1 Marks, H3K27Ac Marks, and transcription factor ChIP-seq data. SLE-associated risk variants (rs58905141, rs10499197) overlap with H3K4Me1 and H3K27Ac Marks and are very close to strong ChIP-seq signals (the darkness of the segment is proportional to the signal strength). The ChIA-PET signal ~20kb upstream of the TNFAIP3 promoter did not overlap with any variants present on the risk haplotype.

### *TNFAIP3* expression is regulated by upstream DNA element

To further analyze the functional potential of the putative enhancer ~ 55kb upstream of *TNFAIP3*, we generated CRISPR-Cas9 constructs to modify the region using the rs10499197 variant as a landmark. We used two sets of small RNAs (guide (g)-RNA) to enable Cas9 mediated genome editing at the target site in HEK293T cells: gRNA#1 was located 31 bp upstream of rs10499197, and gRNA#10 was located 4 bp downstream of rs10499197 (**Supplementary Figure**). Several isogenic clones were Sanger sequenced (**Supplementary Table**), but only three cell clones exhibited varying patterns of CRISPR editing near the target site. Clone#10-14 (from gRNA#10) had a 22-nucleotide deletion around rs10499197 on one of the alleles and 62-nucleotide deletion around rs10499197 on the other (**Figure 2a**). Clone#10-9 had a 11-nucleotide deletion around rs10499197 on one allele and a 9-nucleotide deletion downstream of the target site on the other allele (**Figure 2a**). Clone#1-6 (from gRNA#1) had a homozygous 5-nucleotide deletion upstream of rs10499197 (**Figure 2a**). Western blotting for A20 protein in whole-cell lysates from the CRISPR clones revealed that the homozygous 5-nucleotide deletion upstream of the rs10499197 significantly reduced *TNFAIP3* expression (**Figure 2b**). Our CRISPR technology provided an orthologus approach to support the ENCODE data by demonstrating that the ~ 55 kb region upstream of the *TNFAIP3* promoter functions as a likely enhancer necessary to sustain *TNFAIP3* expression. The data also suggests that the rs10499197 may be a causal variant that disrupts *TNFAIP3* gene expression by disrupting the enhancer.

**Figure 2.**
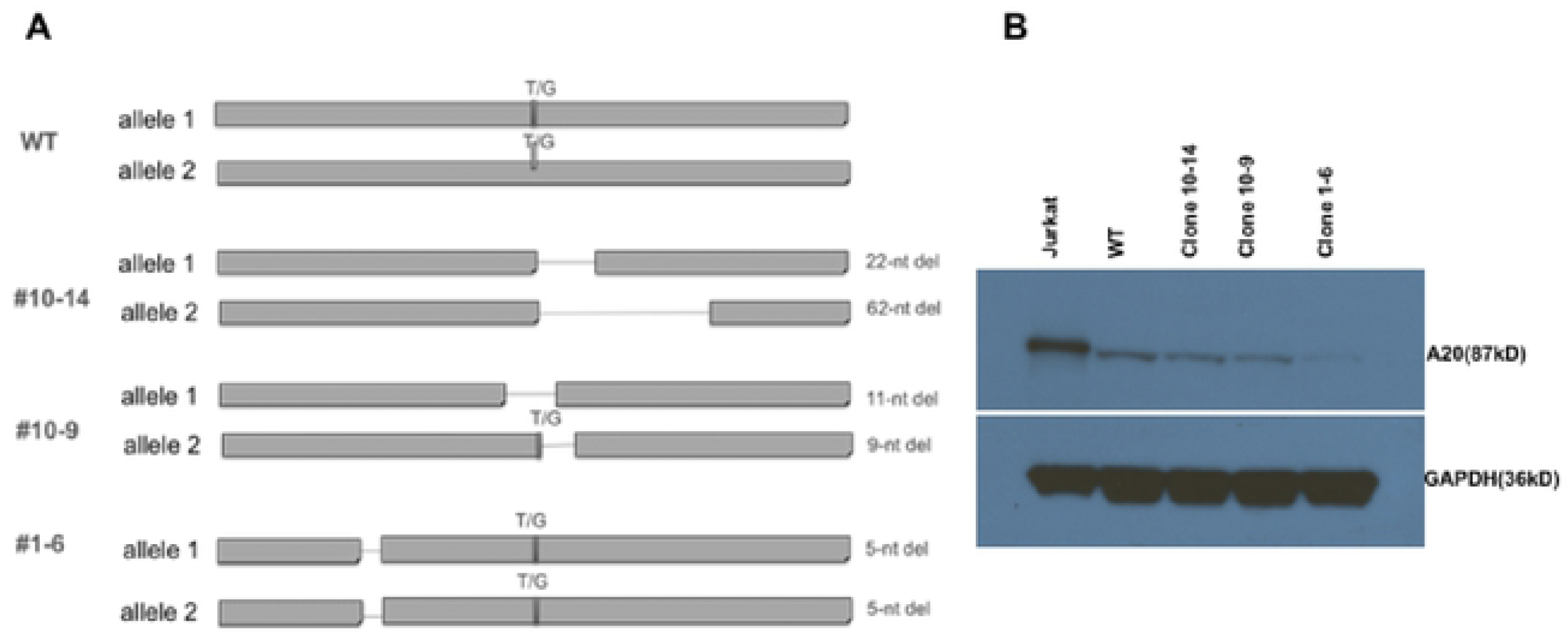
Evaluation of isogenic TNFAIP3 clones following genome editing. (A) illustration showing edited upstream putative enhancer region around rs10499197 across three CRISPR-HEK293T clones: #10-14,10-9, and 1-6. WT represents wildtype cell. The clones either have a homozygous deletion or a heterozygous deletion. (B) A Western blot demonstrating the difference in A20 expression across all the clones when compared to the wildtype. Cell lysate from Jurkat T cells were used as a positive control. GAPDH represents the loading control.

## Discussion

In this study, we describe the functional characterization of the rs10499197 variant that is associated with human SLE in the region of *TNFAIP3* on chromosome 6q23 for which previous genetic and bioinformatics analyses suggest there are likely to be causal variants. A key objective of human genetics is the identification and characterization of variants associated with complex diseases. Although extensive platforms such as GWAS help us identify several variants associated with complex genetic diseases only a small fraction are causal. Due to Linkage Disequilibrium (LD) in the human genome, GWAS is usually unable to differentiate causal variants from non-causal variants on the same haplotype. Therefore, separating causal variants from non-causal variants requires a combination of genetic (finemapping, resequencing, imputation) and bioinformatic (variant annotation, modeling building) techniques. The end result, as a consequence, is a list of variants that methodically characterized to understand the underlying biological function in relation to complex diseases. The current work offers a functional explanation for the genetic association between SLE and the alleles of the variant rs10499197. Thereby making rs10499197 a potential SLE risk haplotype. The *TNFAIP3* locus exhibits complex genetic architecture. A study looking at the genetic associations of SLE and *TNFAIP3* identified three key risk haplotypes around the *TNFAIP3* gene body (Plenge R.M. et al., 2007): 185kb haplotype upstream of *TNFAIP3*, 109kb haplotype that spans the *TNFAIP3* coding region and a 249kb haplotype downstream of *TNFAIP3*. Our work focuses on a non-coding DNA element 55kb upstream of the *TNFAIP3* promoter. This upstream DNA element harbors SLE variants believed to affect *TNFAIP3* and in turn SLE pathophysiology.

## Conclusions

Our future work would include, Chromatin conformation capture assays (3C) that would determine how the upstream DNA element physically interacts with the *TNFAIP3* promoter. An understanding of which will help us establish a novel mechanism for *TNFAIP3* transcriptional regulation.

---

**Supplementary Fig.**
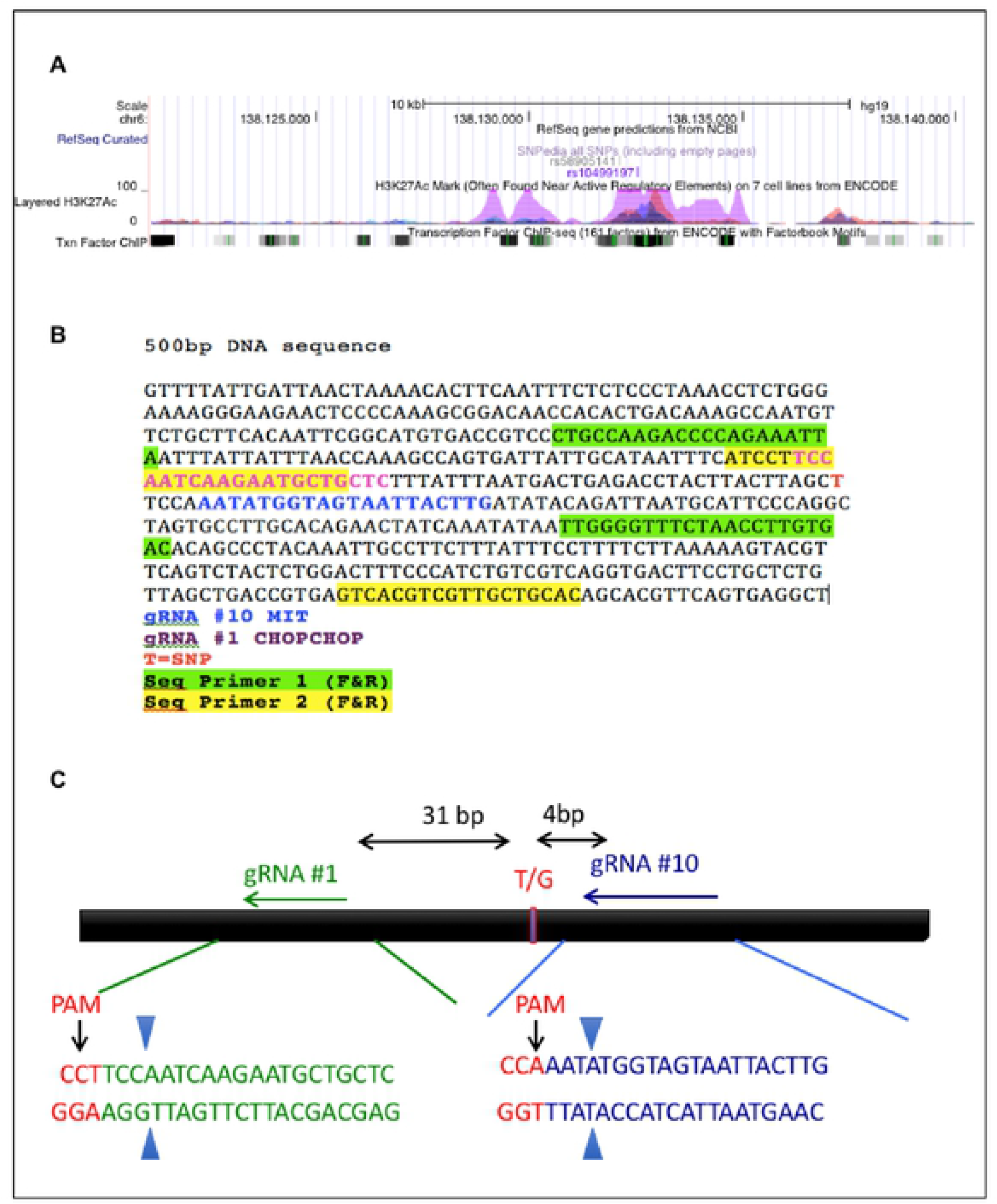
CRISPR-CAS9 guide RNA design. (A) Location of putative enhancer upstream of *TNFAIP3* identified by the risk variant rs10499197 in the UCSC genome browser. The top track shows location of the SNPs, the middle track shows the H3K27ac marks that are indicative of an enhancer region, the bottom track shows the ENCODE defined transcription factor binding tracks (B) Sequence around rs10499197 in the upstream DNA element. (C) location of designed guide RNA, PAM sequence around rs10499197 (T/G).

**Supplementary Table.**
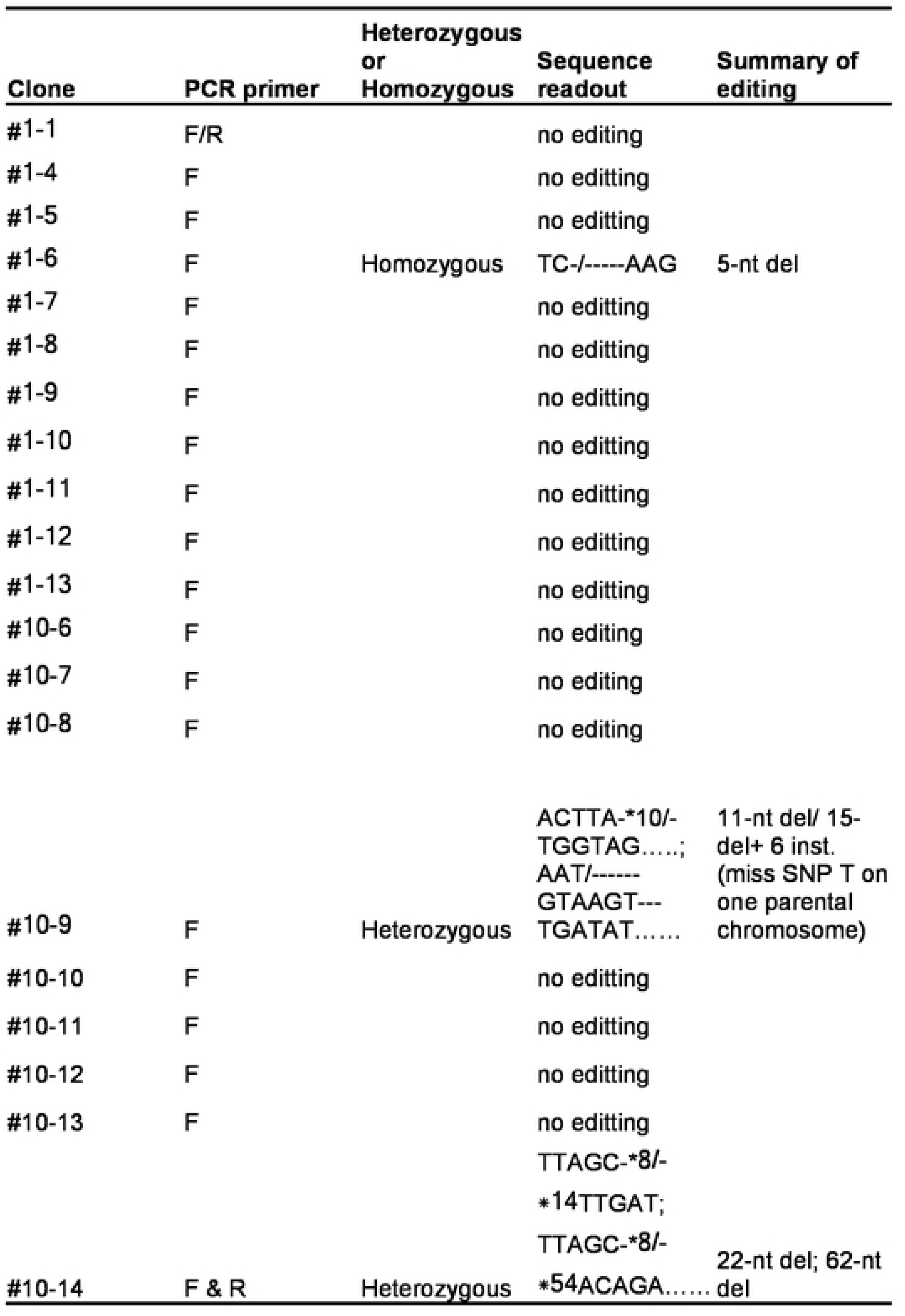
PCR screening of Clones. Table columns show (from left to right): isogenic clones; PCR primers used to sanger sequence the clones; editing in one allele or both; sequence readout of the edit.

## References

Bonev, B., and Cavalli, G. (2016). Organization and function of the 3D genome. Nat Rev Genet 17, 772.

Cai, L.Q., Wang, Z.X., Lu, W.S., Han, J.W., Sun, L.D., Du, W.H., Zhang, S.M., Zuo, X.B., Zhang, X.J., and Yang, S. (2010). A single-nucleotide polymorphism of the TNFAIP3 gene is associated with systemic lupus erythematosus in Chinese Han population. Mol Biol Rep 37, 389–394.

Chen, L., Morris, D.L., and Vyse, T.J. (2017). Genetic advances in systemic lupus erythematosus: an update. Curr Opin Rheumatol 29, 423–433.

Consortium, E.P. (2012). An integrated encyclopedia of DNA elements in the human genome. Nature 489, 57–74.

Das, T., Chen, Z., Hendriks, R.W., and Kool, M. (2018). A20/Tumor Necrosis Factor alpha-Induced Protein 3 in Immune Cells Controls Development of Autoinflammation and Autoimmunity: Lessons from Mouse Models. Front Immunol 9, 104.

Dekker, J. (2006). The three ‘C’ s of chromosome conformation capture: controls, controls, controls. Nat Methods 3, 17–21.

Deng, Y., and Tsao, B.P. (2017). Updates in Lupus Genetics. Curr Rheumatol Rep 19, 68.

Dieude, P., Guedj, M., Wipff, J., Ruiz, B., Riemekasten, G., Matucci-Cerinic, M., Melchers, I., Hachulla, E., Airo, P., Diot, E., Hunzelmann, N., Cabane, J., Mouthon, L., Cracowski, J.L., Riccieri, V., Distler, J., Meyer, O., Kahan, A., Boileau, C., and Allanore, Y. (2010). Association of the TNFAIP3 rs5029939 variant with systemic sclerosis in the European Caucasian population. Ann Rheum Dis 69, 1958–1964.

Genetic Analysis of Psoriasis, C., The Wellcome Trust Case Control, C., Strange, A., Capon, F., Spencer, C.C., Knight, J., Weale, M.E., Allen, M.H., Barton, A., Band, G., Bellenguez, C., Bergboer, J.G., Blackwell, J.M., Bramon, E., Bumpstead, S.J., Casas, J.P., Cork, M.J., Corvin, A., Deloukas, P., Dilthey, A., Duncanson, A., Edkins, S., Estivill, X., Fitzgerald, O., Freeman, C., Giardina, E., Gray, E., Hofer, A., Huffmeier, U., Hunt, S.E., Irvine, A.D., Jankowski, J., Kirby, B., Langford, C., Lascorz, J., Leman, J., Leslie, S., Mallbris, L., Markus, H.S., Mathew, C.G., Mclean, W.H., Mcmanus, R., Mossner, R., Moutsianas, L., Naluai, A.T., Nestle, F.O., Novelli, G., Onoufriadis, A., Palmer, C.N., Perricone, C., Pirinen, M., Plomin, R., Potter, S.C., Pujol, R.M., Rautanen, A., Riveira-Munoz, E., Ryan, A.W., Salmhofer, W., Samuelsson, L., Sawcer, S.J., Schalkwijk, J., Smith, C.H., Stahle, M., Su, Z., Tazi-Ahnini, R., Traupe, H., Viswanathan, A.C., Warren, R.B., Weger, W., Wolk, K., Wood, N., Worthington, J., Young, H.S., Zeeuwen, P.L., Hayday, A., Burden, A.D., Griffiths, C.E., Kere, J., Reis, A., Mcvean, G., Evans, D.M., Brown, M.A., Barker, J.N., Peltonen, L., Donnelly, P., and Trembath, R.C. (2010). A genome-wide association study identifies new psoriasis susceptibility loci and an interaction between HLA-C and ERAP1. Nat Genet 42, 985–990.

Han, J.W., Zheng, H.F., Cui, Y., Sun, L.D., Ye, D.Q., Hu, Z., Xu, J.H., Cai, Z.M., Huang, W., Zhao, G.P., Xie, H.F., Fang, H., Lu, Q.J., Xu, J.H., Li, X.P., Pan, Y.F., Deng, D.Q., Zeng, F.Q., Ye, Z.Z., Zhang, X.Y., Wang, Q.W., Hao, F., Ma, L., Zuo, X.B., Zhou, F.S., Du, W.H., Cheng, Y.L., Yang, J.Q., Shen, S.K., Li, J., Sheng, Y.J., Zuo, X.X., Zhu, W.F., Gao, F., Zhang, P.L., Guo, Q., Li, B., Gao, M., Xiao, F.L., Quan, C., Zhang, C., Zhang, Z., Zhu, K.J., Li, Y., Hu, D.Y., Lu, W.S., Huang, J.L., Liu, S.X., Li, H., Ren, Y.Q., Wang, Z.X., Yang, C.J., Wang, P.G., Zhou, W.M., Lv, Y.M., Zhang, A.P., Zhang, S.Q., Lin, D., Li, Y., Low, H.Q., Shen, M., Zhai, Z.F., Wang, Y., Zhang, F.Y., Yang, S., Liu, J.J., and Zhang, X.J. (2009). Genome-wide association study in a Chinese Han population identifies nine new susceptibility loci for systemic lupus erythematosus. Nat Genet 41, 1234–1237.

Heinz, S., Romanoski, C.E., Benner, C., and Glass, C.K. (2015). The selection and function of cell type-specific enhancers. Nat Rev Mol Cell Biol 16, 144–154.

Lareau, C.A., and Aryee, M.J. (2018). hichipper: a preprocessing pipeline for calling DNA loops from HiChIP data. Nat Methods 15, 155–156.

Mohan, C., and Putterman, C. (2015). Genetics and pathogenesis of systemic lupus erythematosus and lupus nephritis. Nat Rev Nephrol 11, 329–341.

Moulton, V.R., Suarez-Fueyo, A., Meidan, E., Li, H., Mizui, M., and Tsokos, G.C. (2017). Pathogenesis of Human Systemic Lupus Erythematosus: A Cellular Perspective. Trends Mol Med 23, 615–635.

Mumbach, M.R., Rubin, A.J., Flynn, R.A., Dai, C., Khavari, P.A., Greenleaf, W.J., and Chang, H.Y. (2016). HiChIP: efficient and sensitive analysis of protein-directed genome architecture. Nat Methods 13, 919–922.

Plenge, R.M., Cotsapas, C., Davies, L., Price, A.L., De Bakker, P.I., Maller, J., Pe’er, I., Burtt, N.P., Blumenstiel, B., Defelice, M., Parkin, M., Barry, R., Winslow, W., Healy, C., Graham, R.R., Neale, B.M., Izmailova, E., Roubenoff, R., Parker, A.N., Glass, R., Karlson, E.W., Maher, N., Hafler, D.A., Lee, D.M., Seldin, M.F., Remmers, E.F., Lee, A.T., Padyukov, L., Alfredsson, L., Coblyn, J., Weinblatt, M.E., Gabriel, S.B., Purcell, S., Klareskog, L., Gregersen, P.K., Shadick, N.A., Daly, M.J., and Altshuler, D. (2007). Two independent alleles at 6q23 associated with risk of rheumatoid arthritis. Nat Genet 39, 1477–1482.

Schaffner, W. (2015). Enhancers, enhancers - from their discovery to today’s universe of transcription enhancers. Biol Chem 396, 311–327.

Servant, N., Varoquaux, N., Lajoie, B.R., Viara, E., Chen, C.J., Vert, J.P., Heard, E., Dekker, J., and Barillot, E. (2015). HiC-Pro: an optimized and flexible pipeline for Hi-C data processing. Genome Biol 16, 259.

